# Development of OPLS-AA/M Parameters for Simulations of G Protein-Coupled Receptors and Other Membrane Proteins

**DOI:** 10.1101/2022.01.05.475148

**Authors:** Michael J. Robertson, Georgios Skiniotis

## Abstract

G protein-coupled receptors (GPCRs) and other membrane proteins are valuable drug targets, and their dynamic nature makes them attractive systems for study with molecular dynamics simulations and free energy approaches. Here, we report the development, implementation, and validation of OPLS-AA/M force field parameters to enable simulations of these systems. These efforts include the introduction of post-translational modifications including lipidations and phosphorylation. We also modify previously reported parameters for lipids to be more consistent with the OPLS-AA force field standard and extend their coverage. These new parameters are validated on a variety of test systems, with the results compared to high-level quantum mechanics calculations, experimental data, and simulations with CHARMM36m where relevant. The results demonstrate that the new parameters reliably reproduce the behavior of membrane protein systems.

## Introduction

G protein-coupled receptors are one of the most important classes of drug targets, comprising roughly 30-40% of modern pharmaceuticals^1^. All GPCR families possess a seven transmembrane helix domain but differ substantially in their extracellular domains and activation mechanisms. However, despite similar overall architecture, members of the same GPCR family can also display incredible diversity in ligand binding domains; family A, for example, contains members that couple to small molecules, peptides, lipids, glycoproteins, photons, and protons. The intracellular ends of each receptor provide selectivity for a distinct subset of G protein subunits that control which secondary messenger(s) signaling cascade is initiated^2^. In addition to G proteins, the intracellular side of GPCRs also couple to the arrestins in order to desensitize and regulate GPCR signaling^3^ as well as mediate distinct signaling pathways^4^. The propagation of activation between the extracellular ligand binding domain and the intracellular domain is a complicated and dynamic process, with experimental evidence for a complex energy landscape with some receptors occupying several distinct intermediate conformations^5^. Further, GPCRs exhibit a range of interactions with ligands, including inverse agonism where a ligand stabilizes the inactive state of a receptor and reduces constitutive activity; neutral antagonism, where there is no influence on receptor activation state but other modulators are unable to bind; and partial and full agonism, where a receptor is induced into an active state, either partially or fully^6,7^. Further, some agonists are capable of stabilizing a receptor in distinct conformations that exhibit preferential coupling with different signaling partners compared to other agonists of the same receptor^2^. There are thus numerous aspects of GPCRs and their signaling that are ideal candidates for study with computational biophysics simulations. However, performing simulations with GPCRs and their interaction partners can be particularly challenging given the wide range of nonstandard biomolecular parameters required for their simulations.

As GPCRs are membrane proteins, accurate simulations require well-validated parameters for a variety of model neutral and anionic lipids as well as crucial physiological lipids like cholesterol and PIP2, as lipid composition plays key roles in GPCR localization and function^8^ (Figure 1). PIP2, for example, binds to arrestin and stabilizes its interaction with both the cellular membrane and with GPCRs upon coupling (Figure 1c)^9,10^. GPCRs and their effectors are also targets for a variety of post-translational modifications. Lipidation on the C-terminus/helix 8^11^ (Figure 1a) is prevalent and confers membrane anchoring that plays varied roles in receptor stability, function, and localization^12–15^. Phosphorylation on the C-terminus and intercellular loop 3 of GPCRs is responsible for receptor desensitization and internalization by arrestins^16,17^ (Figure 1b). The G protein heterotrimer also bears multiple lipidations sites to ensure membrane anchoring^12^ (Figure 1d). Both of these families of post-translational modifications and the phospholipid bilayer itself need to be accurately characterized in a force field in order to perform realistic simulations of GPCR systems.

**Figure 1.**
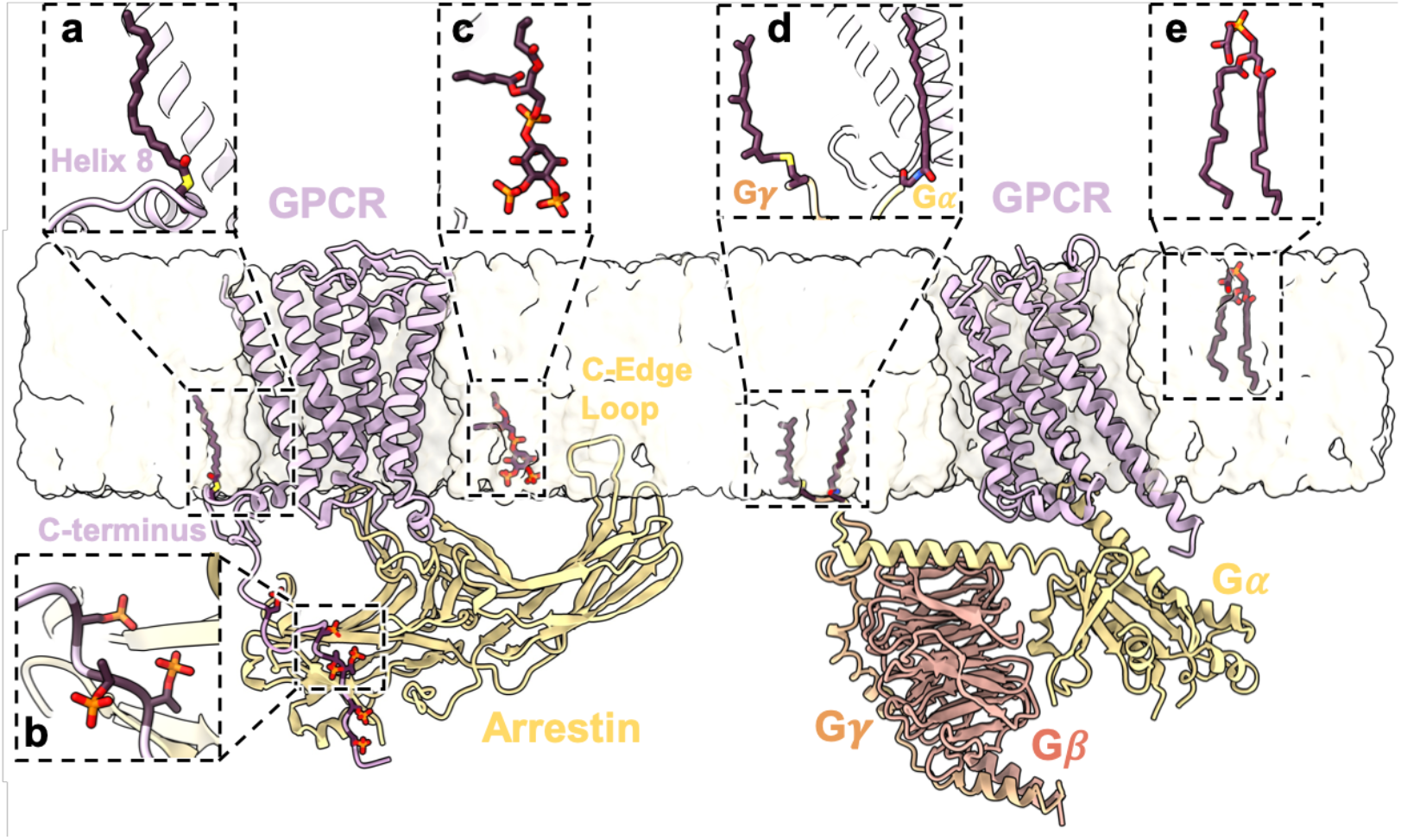
Examples of components parameterized in this work (a) Cysteine palmitoylation on helix 8 of a GPCR. (b) Phosphorylation of a GPCR C-terminal tail at multiple serine and threonine sites. (c) PIP2 interaction site for an arrestin-GPCR complex (d) Cysteine prenylation of Gγ and N-terminal glycine myristoylation of Gαi. (e) POPG lipid bilayer component

There are several popular choices of biomolecular force fields, including CHARMM^18^, AMBER, GROMOS, and OPLS-AA^19^. The first three of these have published parameters for a full suite of lipids and post-translational modifications. The coverage of the CHARMM force field is particularly extensive and is complemented by the CHARMM-GUI^20^ to assist in building PDB files for a wide variety of systems. The OPLS-AA force field has recently received modern parameter updates and validation for its major biomolecular components, including proteins^21^ and RNA^22^. However, in contrast to other force fields, parameterization beyond very standard biomolecular components has been limited to date.

Here, we report the parameterization and expansion of the OPLS-AA/M force field to allow for GPCR simulations with and without their effectors to be performed. This includes development of new amino acid χ_1_ dihedral parameters and thioester nonbonded parameters for the addition of N- and S-palmitoylation, and the introduction of parameter and topology files for N-myristoylation and S-farnesylation to cover lipid anchoring of GPCRs and G proteins. Given the substantial importance of GPCR phosphorylation in arrestin coupling and signaling, we also develop parameters for di- and mono-basic phosphorylation of serine and threonine. While parameters have been reported previously for OPLS-AA phospholipids and cholestrol^23^, their GROMACS-based format does not fully align with the OPLS-AA/M parameterization philosophy, and thus new torsion terms have been derived to conform. Finally, the new systems are validated on a variety of test cases, including molecular dynamics of a muscarinic 2 (M2) receptor/β-arrestin complex and free energy perturbation simulations with the well-studied cannabinoid 1 (CB1) receptor, with the results comparing favorably to both experiment and the CHARMM36m force field. All parameter and topology files generated in this work are provided in the CHARMM format to allow the membrane-system building utilities of the CHARMM-GUI to be leveraged in simulation setup.

## Results

### Phosphorylation

We initially performed molecular dynamics simulations of a series of blocked GSXS peptides (Figure 2a), where X was either serine or threonine in standard, monobasic phosphorylated, or dibasic phosphorylated form. We chose this test system as there is an available NMR study that probes the effect of phosphorylation on the backbone 3J couplings^24^, which have proven an invaluable metric for force field assessment. Based on the experimental results, phosphorylation of the central residue should induce a significant downward shift in certain backbone 3J couplings, as interaction with the charged sidechain induces more alpha-helical character in the conformational ensemble. The existing OPLS-AA/M serine and threonine dihedral torsion parameters performed well for the standard and monobasic amino acids, however the results for dibasic phosphoserine and phosphothreonine were markedly less accurate (Figure 2b, SI Table 1). Consistent with prior parameterization efforts, OPLS-AA/M parameters for dibasic phosphorylated serine χ_1_ were derived from QM scans of a blocked phosphorylated serine dipeptide (SI Figure 1), with the same χ_1_ parameter also used for phosphothreonine. These newly derived parameters substantially improved the agreement with the experimental 3J couplings, dropping from an RMSE of 0.58 Hz to 0.42 Hz for dibasic phosphoserine and 1.27 Hz to 0.38 Hz for dibasic phosphothreonine.

**Figure 2.**
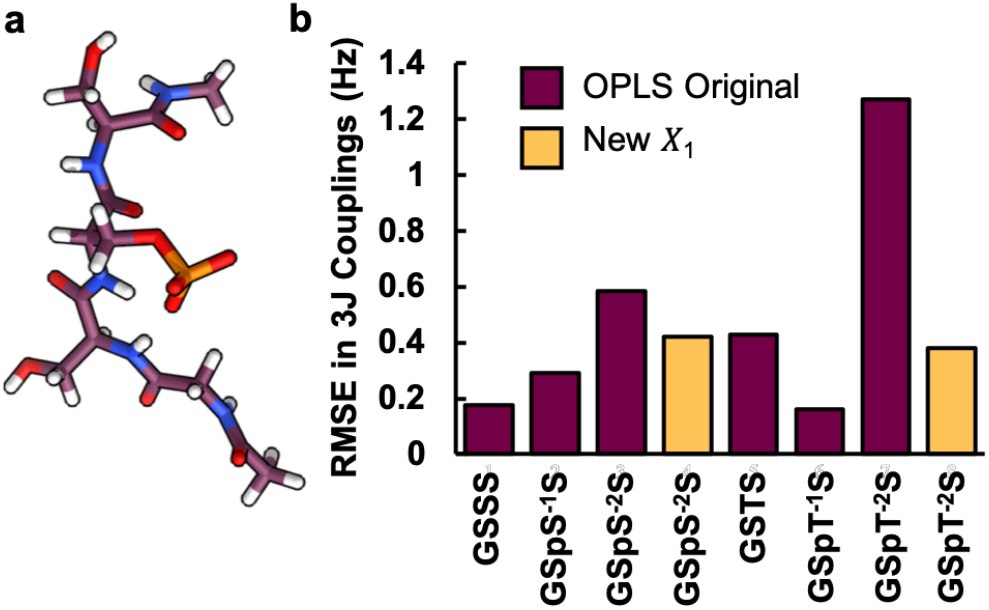
Molecular dynamics results for GSXS peptides (a) Snapshot of GSpS^-2^S peptide from MD simulations. (b) Root mean squared errors in calculated 3J couplings for GSXS peptides.

### Phospholipid Bilayers

Kulig *et al*.^23,25^ previously reported GROMACS-formatted OPLS-AA parameters for several lipids including POPC, DPPC, DOPC and cholesterol. They modified both dihedral and nonbonded terms and obtained excellent agreement for experimental data on phospholipid bilayer properties including area per head group and compressibility. However, while the nonbonded parameters can easily be transferred to CHARMM and OPLS format, dihedral terms in GROMACS are provided in Ryckaert-Bellemans notation (Equation 1) rather than Fourier dihedrals (Equation 2), and conversion to Fourier dihedrals yields V_4_ terms, which are generally reserved for systems where fourfold minima are expected (*e.g*., some biaryl torsions) to prevent overfitting. Thus, we have repeated the scans described in the original paper at the ωB97-xd (6-311+(2d,2p) level (SI Fig 2), with the exception of the phosphate OS-glycerol branching point parameters OS-CT-CT-OS and OS-CT-CT-CT, which were scanned using a construct akin to that of Klauda *et al*.^26^ (SI Fig 2k). We additionally performed scans for phosphoserine, phosphoethanolamine, and phosphoglycerol (PS, PE, and PG) headgroups (SI Fig 2) that were not previously parameterized. The updated parameters were used to simulate pure bilayers of POPC, DPPC, DOPC, POPE, and POPS, with area per headgroup and compressibility performing well compared to both experimental data (Table 1) and values reported in the literature for other popular force fields^20^. Further, topology and parameter entries for cholesterol and PIP2 were prepared by combining our new results with existing parameters from the OPLS-AA force field and Rog *et al*.

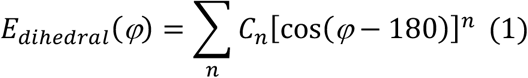

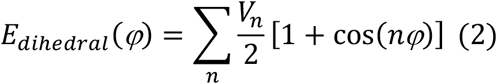

**Table 1:**
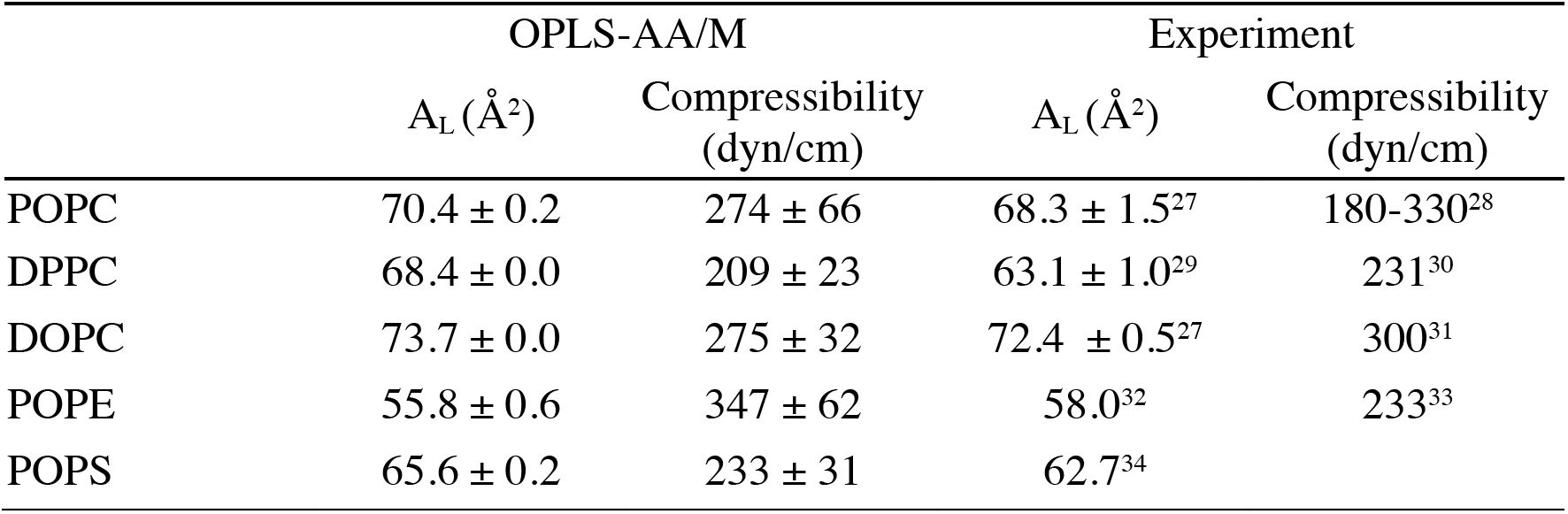
Calculated properties for phospholipid bilayers with OPLS-AA/M parameters.

### Lipidation

N- and S-palmitoylation, N-myristoylation, S-farnesylation, and S-geranylgeranylation are the most common lipidations we chose to parameterize. To incorporate S-palmitoylation, nonbonded parameters for thioesters were developed based upon existing thiol and ester OPLS-AA parameters and several dihedrals were fit to QM scans (SI Fig 3). The results were validated with pure liquid Monte Carlo simulations of simple thioesters, which demonstrated good performance in reproducing experimental densities and heats of vaporization (SI Table 2). We then developed new χ_1_ parameters for S-palmitoylation and S-farnesylation/S-geranylgeranylation based on blocked thioester and sulfide analogues of cysteine as model compounds (SI Fig 4,5). Topology and parameter files for the lipidated cysteines were then prepared by combining these results for peptide and thioester/sulfide linkages with our updated lipid parameters for the hydrocarbon tails. We are unaware of any NMR validation data for small lipidated cysteine peptides, likely due to the complexities of performing quantitative experiments on these peptides. In lieu of this, we compare the behavior of the lipidation in our CB1 validation systems to the CHARMM36m results, as detailed in the following section. No new additional dihedral parameters are necessary for N-terminal lipidations, as amide parameters are well-established in the OPLS-AA force field, so topology and parameter files were again prepared by combination of existing parameters with our new lipid parameters.

### CB1 FEP Calculations

To probe how well our new lipid parameters provide an accurate physical environment for a membrane protein, we performed free energy perturbation simulations on a series of phytocannabinoids bound to cannabinoid receptor 1 (CB1). Cannabinoid receptor 1 is a family A GPCR that is the most widely-expressed GPCR in the brain where it plays a key role in neurotransmitter release from pre-synaptic neurons. It has also been the subject of a great deal of structural characterization and compound development, which provides rich SAR data for validation purposes. CB1 is also C-terminally palmitoylated, crucial for proper receptor membrane localization and signaling^15^. We chose a series of the most salient historical optimizations to a Δ8-tetrahydrocannabinol (THC) based scaffold^35^ (Figure 3a, SI Fig 6) to use for free energy perturbation calculations. Both OPLS-AA/M and CHARMM36m performed well in reproducing the relative free energies of binding, with almost all perturbations having the correct sign (83%, SI Table 3) and with good agreement for calculated free energies of binding from the initial reference compound (MUE in individual ΔΔGs of 1.11 kcal/mol for OPLS-AA/M and 0.94 kcal/mol for CHARMM36m, MUE in ΔG of 0.73 kcal/mol for OPLS-AA/M and 1.21 kcal/mol for CHARMM36m). The most significant ΔG outlier corresponds to the Δ8-THC analogue with a methylated hydroxyl in the C1 position. For both force fields, conversion of the hydroxyl to the ether was strongly disfavored (2.3-2.5 kcal/mol), however experimentally the measured change is even greater at 4.3 kcal/mol. While qualitatively this result is correct, the large quantitative difference suggests either a systematic deficiency in both force fields, an error in the experimental data, or a need for improved sampling, *e.g*., with grand canonical Monte Carlo to better sample water-mediated interactions.

**Figure 3.**
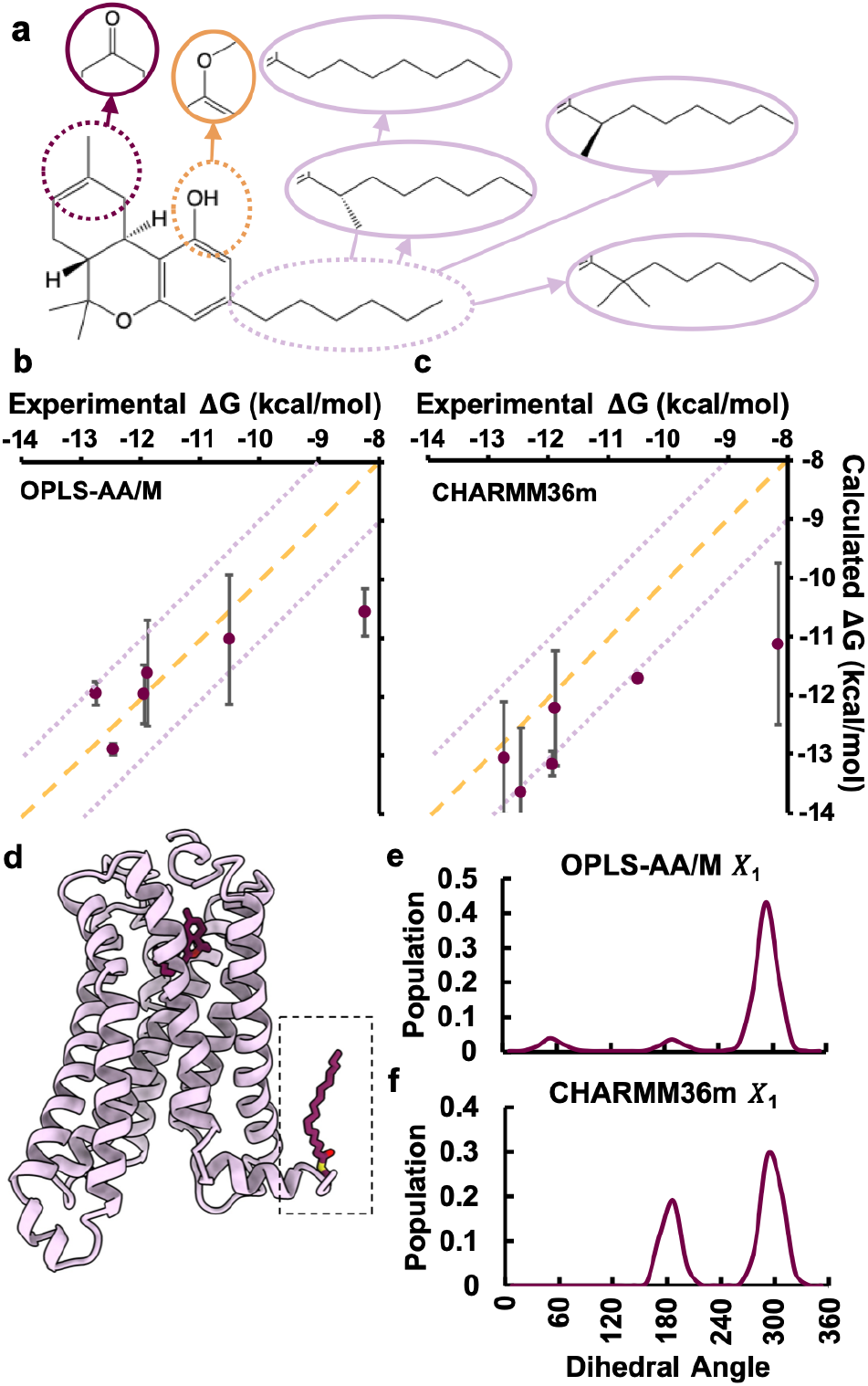
FEP Simulations of Cannabinoid Receptor 1 (a) Diagram of the phytocannabinoid analogues subjected to FEPs. (b,c) Plots of experimental vs calculated free energies of binding calculated relative to the initial compound with the OPLS-AA/M (b) and CHARMM36m (c) force fields. (d) Cartoon of CB1 showing lipidation of helix 8 and the bound phytocannabinoid ligand. (e,f) Dihedral populations of palmitoylated cysteine χ_1_ from CB1 simulations with OPLS-AA/M (e) and CHARMM36m (f).

All simulations of CB1 were performed in a POPC bilayer with palmitoylation on the helix 8 cysteine, utilizing both our new OPLS-AA/M parameters and CHARMM36m, providing the opportunity to compare the performance of our new sidechain dihedral parameters for palmitoylation. The palmitoylation cysteine χ_1_ predominantly occupied the ‘m’ (N-Cα-Cβ-Cγ = −60°) rotamer with both force fields. OPLS-AA/M had equal minor populations of the ‘p’ (N-Cα-Cβ-Cγ = 60°) and ‘t’ (N-Cα-Cβ-Cγ = 180°) rotamers, while CHARMM36m never occupied the p rotamer. There are an insufficient number of experimental structures with well-resolved palmitoylation sites to compare these results to a PDB survey, however these results are in line with expected rotamer populations from cysteine and amino acids in general^36^.

### M2 Arrestin Simulations

As a final validation, we performed simulations of phosphorylated muscarinic acetylcholine receptor 2 (M2R)/β-arrestin 1 complex in either a small lipid nanodisc or a POPC/POPG bilayer, a pair of simulations described previously^37^, with our updated parameters. The full interaction of M2R and arrestin depends on the arrestin C edge loop region (Figure 1) embedding into a membrane environment^37^. Simulations of the complex in a nanodisc that is insufficiently large for the C edge loop to be buried in the phospholipid bilayer results in the arrestin adopting a conformation more like that in the ‘inactive’ state of arrestin^37^ that is less primed for coupling to the phosphorylated receptor, as measured by the interdomain angle between the two lobes of the arrestin^38^. In our new simulations, we recapitulate this effect in good agreement with our prior study (SI Figure 7), again demonstrating that in the absence of a full lipid bilayer arrestin rapidly shifts to a more inactive-like domain arrangement.

This simulation also provides the opportunity to examine the rotamer populations of phosphorylated serine and threonine in the phosphopeptide region of the receptor embedded in the cleft of arrestin (Fig 4d). By comparing our simulated rotamer distributions to the χ_1_ value from the cryoEM and crystal structures of vasopressin phosphopeptide-bound arrestin (Fig 4a-c, e-g; dashed lines) we can see there is good agreement between the simulated value and the structure, with the local protein environment producing correct rotamer population shifts. Further, the phosphoserine residues at the more loosely bound ends of the peptide sample additional conformations, particularly phosphoserine 495 which is in part coordinated by an antibody fab fragment in the experimental structures that is removed for the simulations.

**Figure 4.**
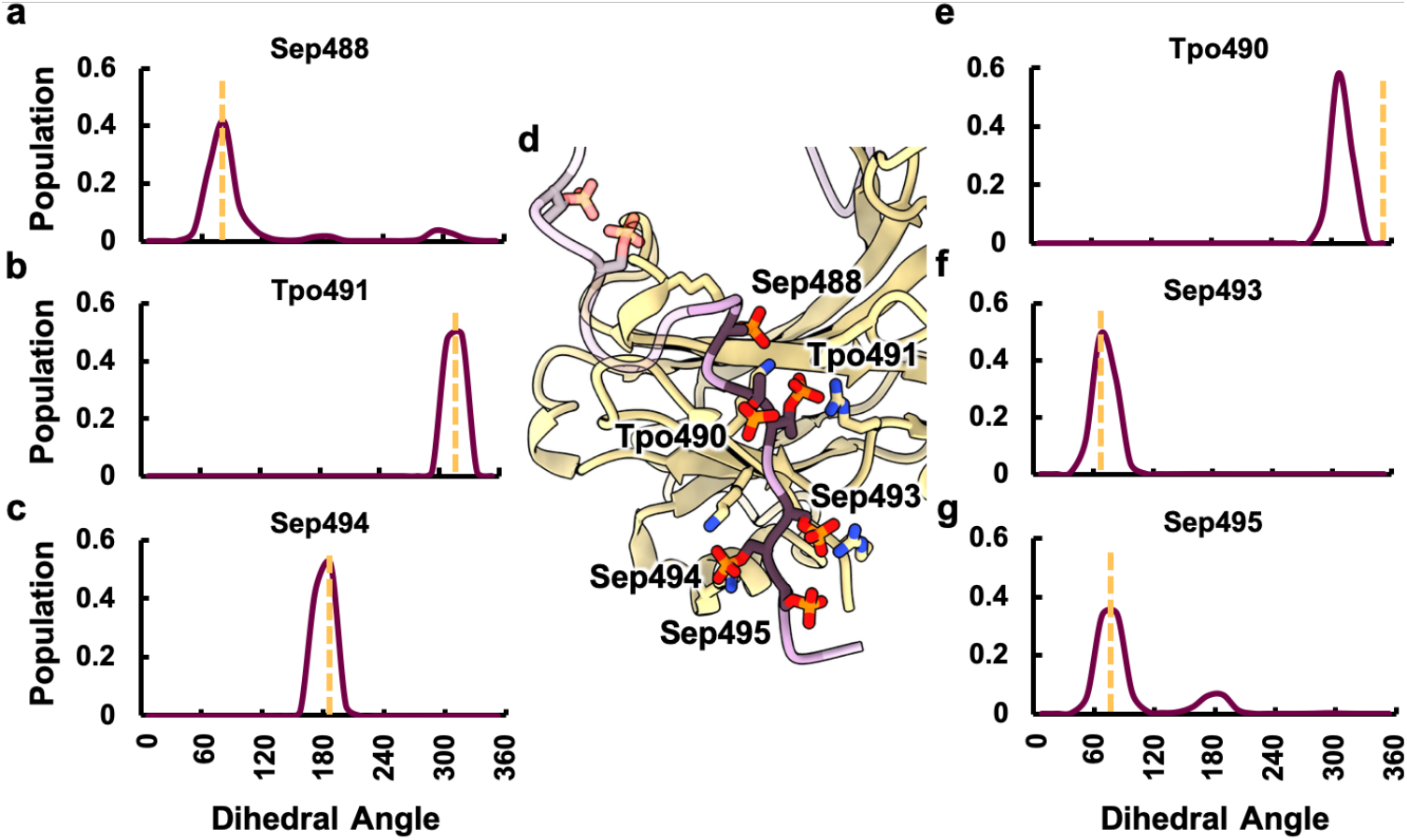
Molecular dynamics simulations of the M2-arrestin complex in a lipid bilayer. (a-c, e-g) Plots of χ_1_ dihedral angle distributions for phosphoserine and phosphothreonine residues in the phosphorylated C-terminus of M2 receptor. Yellow dashed lines correspond to the values from the experimental structure. (d) Snapshot of the arrestin-phospholipid complex from MD simulations with the measured phosphorylated residues labeled. Residues in the C-terminus modeled in the simulation but absent in the structure are transparent.

## Conclusion

We have extended the coverage of the OPLS-AA/M force field to allow for high accuracy simulations of membrane systems and many post-translationally modified proteins. While some unparameterized areas remain, perhaps most notably glycosylation, these new parameters open up a vast array of complicated biochemical systems to study with the OPLS-AA/M force field. Further, many of the validation systems presented here are quite novel and should find further use in continuing force field development. By providing our new parameter and topology files in the CHARMM format, they can be easily applied to membrane and lipid nanodisc systems built with the CHARMM-GUI^20^, greatly simplifying the process of generating input coordinate files. Together with OPLS-AA/M parameters for proteins^21^ and RNA^22,39^, the OPLS-AA/CM1A force field for small molecules^40^, and available tools for automatic OPLS-AA ligand parameterization^41^, this work thus provides the final missing piece for easy and accurate biophysical characterization and structure based drug discovery for GPCRs, ion channels, cytokine receptors, and other membrane proteins of vital importance to human health with the OPLS-AA family of force fields.

## Methods

### Quantum Chemistry & Parameter Fitting

Quantum chemical scans were performed at the ωB97X-D/6-311+G(2d,2p) level of theory in the Gaussian 16 software^42^. All dihedrals were scanned in 15° increments with most other heavy atom dihedrals held fixed. Molecular mechanics scans of the same molecules were performed in an equivalent manner utilizing the BOSS software^43^ and the OPLS-AA force field for all other force field terms not being fit. Dihedral terms were fit as described previously^21,22^, including the usage of a Boltzmann weighting factor with a temperature of 2000 K, although this generally only had an effect on the parameter fits for the peptide side chain dihedrals. For peptide sidechains, blocked dipeptides were scanned in either alpha helix, beta sheet, or polyproline II helix conformations with χ_2_ held in either gauche+, gauche-, or trans configuration and parameters were fit to simultaneously optimize all 9 scans.

### Molecular Dynamics System Preparation

Solvated and ionized coordinate files for molecular dynamics simulations involving a phospholipid bilayer were prepared using the CHARMM-GUI^20^ while those in aqueous phase were prepared using VMD^44^. Initial coordinates for the CB1 receptor-Δ8 THC analogue were built from PDB:5XR8^45^ bound to a closely analogous phytocannabinoid compound with ICL3 manually rebuilt in a loop conformation to replace the lysozyme fusion, palmitoylation on C415, and residues D163 and D213 were protonated as aspartate residues at this position in family A GPCRs are typically protonated when the receptor is in the active conformation. CB1R was simulated in a mixed POPC/CHS bilayer system, with the specific cholesterol modeled in the experimental structure retained. Simulations of M2R/β-Arrestin started from the same receptor-arrestin complex structure preparation described previously^37^. All coordinate files were prepared for molecular dynamics simulation in VMD^44^ to generate PSF files for simulation in NAMD^46^. Ligand parameter files with the CHARMM36m force field were generated with the paramchem webserver for CGenFF parameters^47,48^. OPLS-AA ligands were simulated with OPLS-AA/CM1A^40^ parameters. Protein components used either CHARMM36m or OPLSAA/M^21,49^.

### Molecular Dynamics Simulations

All simulations were executed in NAMD with a constant temperature and pressure of 1 atm maintained using a Nose-Hoover Langevin piston barostat with a piston period of 150 fs and a piston dampening time scale of 75 fs and a Langevin thermostat with a damping coefficient of 1 ps^-1^. Simulations in aqueous phase were performed at 283K, while for most membrane simulations, a temperature of 303.15K was employed, with the exception of POPS (300K), POPE (310K), DOPC (310K), and DPPC (323K). Nonbonded cutoffs were employed at 11 Å, with smoothing at 9 Å and particle mesh Ewald used for long-range electrostatics. A 2 fs time step was employed with the use of SHAKE and SETTLE^50^.

All systems were subjected to 1500 steps of minimization and gradual heating from 0K to the final temperature in 20K intervals with 0.4 ns of simulation at each interval. For simulations of phosphopeptides, triplicate 500 ns were performed. In the case of simulations of model bilayers alone, triplicate 300 ns were performed with the first 100 ns discarded as equilibration. For simulations of CB1, the system was simulated in triplicate forward and backward by being subjected to an additional 10 ns of equilibration at 303.15 K with harmonic restraints of 1.0 kcal/mol/Å^2^ on all non-water, non-ion, non-hydrogen atoms; 5 ns of equilibration with harmonic restraints of 1.0 kcal/mol/Å^2^ on all non-hydrogen protein atoms; followed by 6 ns of simulation with restraints on all CA atoms in the protein slowly stepped down from 1 kcal/mol/Å^2^ to 0 kcal/mol/Å^2^; before FEP simulations with a lambda schedule in increments of 0.025 for windows between 0.0-0.1 and 0.9-1.0 and increments of 0.050 between 0.1-0.9, with each window simulated for 1 ns of equilibration and 5 ns of production. For the M2R arrestin simulation, the system was simulated in five replicates in a lipid nanodisc and a membrane by being subjected to an additional 10 ns of equilibration at 303.15 K with harmonic restraints of 1.0 kcal/mol/Å^2^ on all non-water, non-ion, non-hydrogen atoms; 10 ns of equilibration with harmonic restraints of 1.0 kcal/mol/Å^2^ on all non-hydrogen protein atoms; followed by 10 ns of simulation with restraints on all CA atoms in the protein slowly stepped down from 1 kcal/mol/Å^2^ to 0 kcal/mol/Å^2^, before being simulated for an additional 30 ns of equilibration and 200 ns of production. Monte Carlo simulations of pure liquid thioesters were performed with the BOSS^43^ software package with 5 million steps of equilibration and 50 million steps of production.

### System Analysis

Backbone 3J coupling values for GSXS peptides were calculated as described previously^21^ using the Karplus parameters of Hu and Bax^51^. For the phospholipid bilayer simulations, area per headgroup is calculated by dividing the system area in the XY plane by the number of lipids in a leaflet, and compressibility is calculated with the formula described by Lee *et al*.^20^. CB1R simulations were processed in VMD ParseFEP^52^, where forward and backward FEP simulations were combined with the Bennett acceptance-ratio (BAR) estimator, and error bars reported are the standard deviation over triplicate combined simulations. Arrestin interdomain twist angle for arrestin-M2 simulations was calculated with the formula of Latorracca *et al*., as described previously^37,38^.

## Supporting information

All supplemental tables and figures

## ASSOCIATED CONTENT

### Supporting Information

The following files are available free of charge.

Computational details of QM scans and MM fitting, Monte Carlo simulation results, FEP & MD results (PDF)

All parameter and topology files in the CHARMM format are provided at https://github.com/mjrober101

## AUTHOR INFORMATION

### Author Contributions

The manuscript was written through contributions of all authors. All authors have given approval to the final version of the manuscript.

## ACKNOWLEDGMENT

We would like to acknowledge the Extreme Science and Engineering Discovery Environment (XSEDE)^53^ resource comet-gpu through sdsc-comet allocation TG-MCB190153 (G.S.), which is supported by National Science Foundation grant number ACI-1548562, and the Mathers Foundation (G.S.).

## ABBREVIATIONS

CB1: cannabinoid receptor 1
GPCR: G protein-coupled receptor
M2: muscarinic acetylcholine receptor 2
FEP: free energy perturbation

