## Supplementary material for "Development of OPLS-AA/M Parameters for Simulations of G Protein-Coupled Receptors and Other Membrane Proteins": All supplemental tables and figures

**Table S1:** 3J Couplings for GSXS phosphopeptides.

| GSXS, X= | | Residue 2 3J (Hz) | Residue 3 3J (Hz) | Residue 4 3J (Hz) |
|---|---|---|---|---|
| Ser | Experiment | 6.78 | 6.88 | 7.17 |
| | OPLS-AA/M | 6.99 | 7.10 | 7.19 |
| $pSer^{-2}$ | Experiment | 6.65 | 5.48 | 6.93 |
| | OPLS-AA/M Serine | 7.29 | 6.26 | 6.99 |
| | OPLS-AA/M New | 7.30 | 5.68 | 7.20 |
| $pSer^{-1}$ | Experiment | 6.66 | 6.53 | 7.1 |
| | OPLS-AA/M Serine | 7.02 | 6.21 | 7.25 |
| Thr | Experiment | 6.67 | 7.82 | 7.03 |
| | OPLS-AA/M | 6.96 | 7.14 | 7.11 |
| $pThr^{-2}$ | Experiment | 6.98 | 5.41 | 7.11 |
| | OPLS-AA/M Serine | 7.13 | 7.39 | 6.16 |
| | OPLS-AA/M New | 7.19 | 5.95 | 6.80 |
| $pThr^{-1}$ | Experiment | 6.93 | 7.29 | 7.03 |
| | OPLS-AA/M Serine | 7.08 | 7.11 | 6.88 |

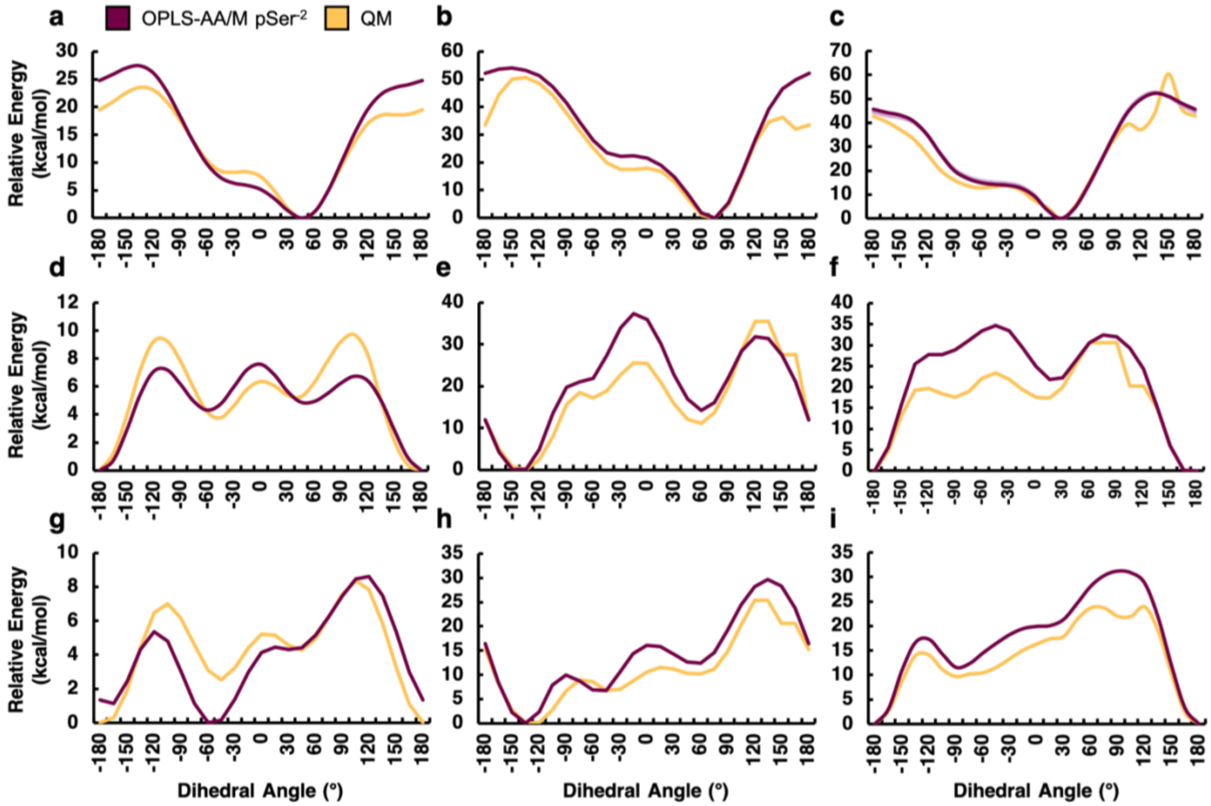


**Figure S1.** Comparison of quantum chemical scans and new OPLS-AA/M force field fits for dibasic phosphoserine in the alpha helical (a,b,c; $\chi_2$= 180, -60, 60), beta sheet (d,e,f; $\chi_2$= 180, -60, 60), and polyproline II (g,h,i; $\chi_2$= 180, -60, 60) conformations.

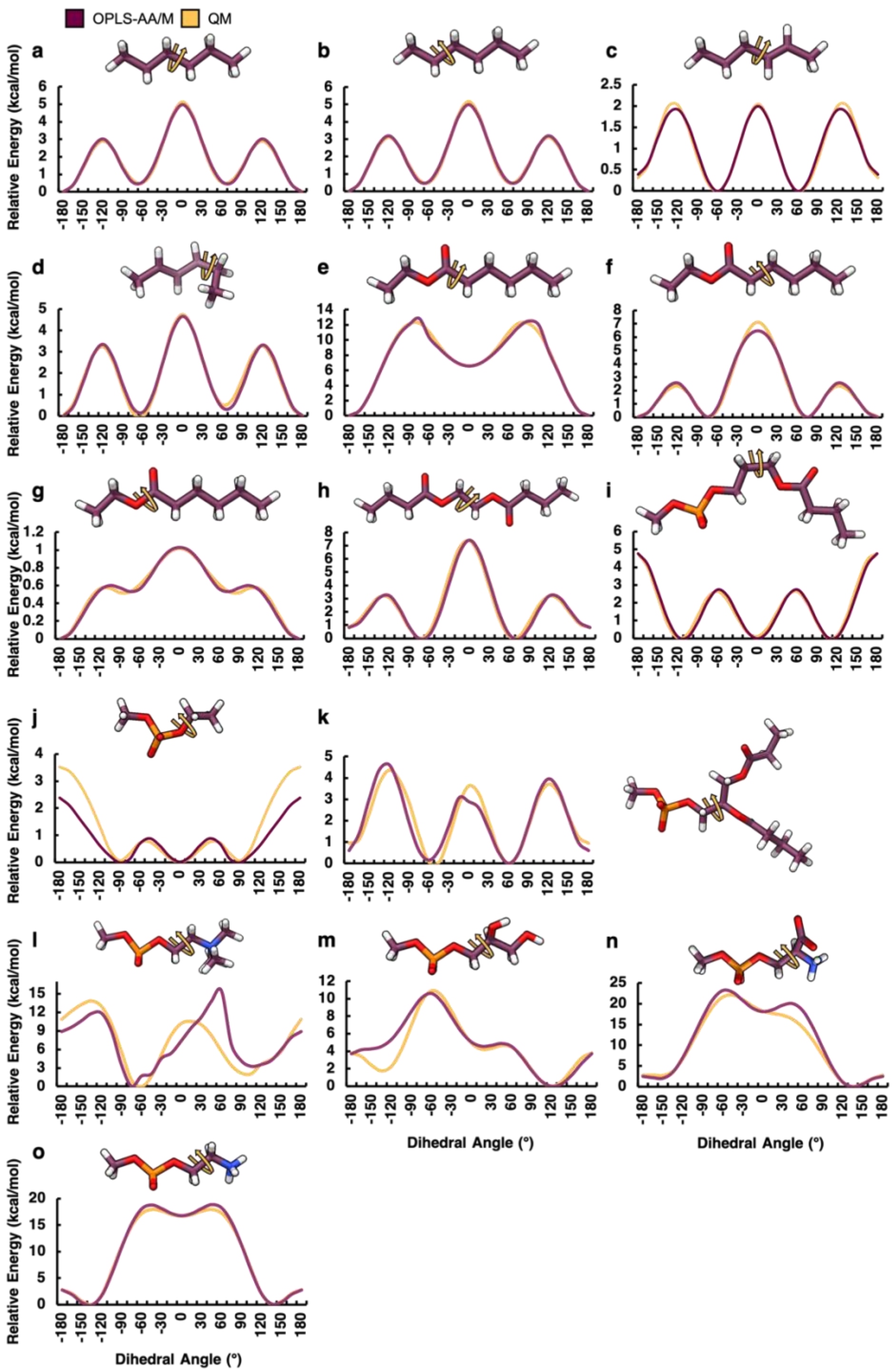


**Figure S2.** Comparison of quantum chemical scans and new OPLS-AA/M force field fits for various phospholipid analogue molecules; hexane (a,b), 2-hexene (c,d), ethyl hexanoate (e-g), ethylene glycol dibutyrate (h), propyl butanoate methyl phosphate (i), ethyl methyl phosphate (j), 1,2-diethanoyl phosphatidyl (k), methyl phosphocholine (PC, l), and methyl phosphoglycerol (PG, m), methyl phosphoserine (PS, n), methyl phosphoethanolamine (PE, o).

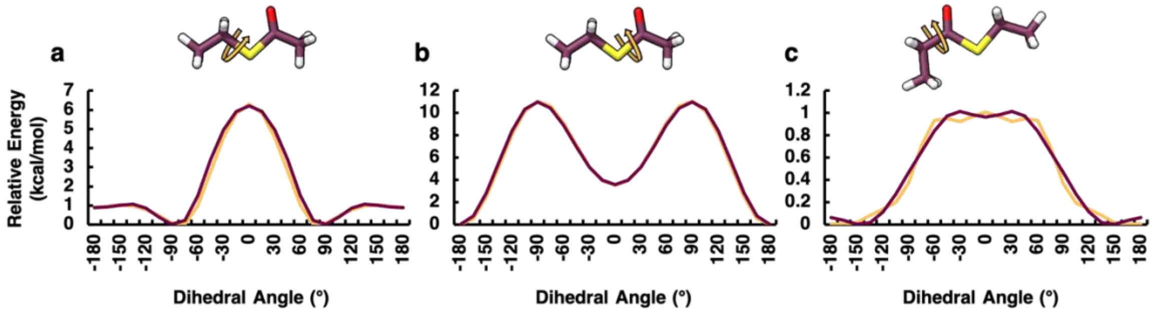


**Figure S3.** Comparison of quantum chemical scans and new OPLS-AA/M force field fits for thioesters.

**Table S2:** Pure liquid properties for thioesters parameterized in this work

| | Calculated | | Experimental | |
|---|---|---|---|---|
| | Density (g/cm$^3$) | $\Delta H_{vap}$ (kcal/mol) | Density (g/cm$^3$) | $\Delta H_{vap}$ (kcal/mol) |
| S-methyl thioate | 1.024 | 8.6 | 1.029 | |
| S-ethyl thioate | 0.979 | 9.6 | 0.991 | 10.4 |

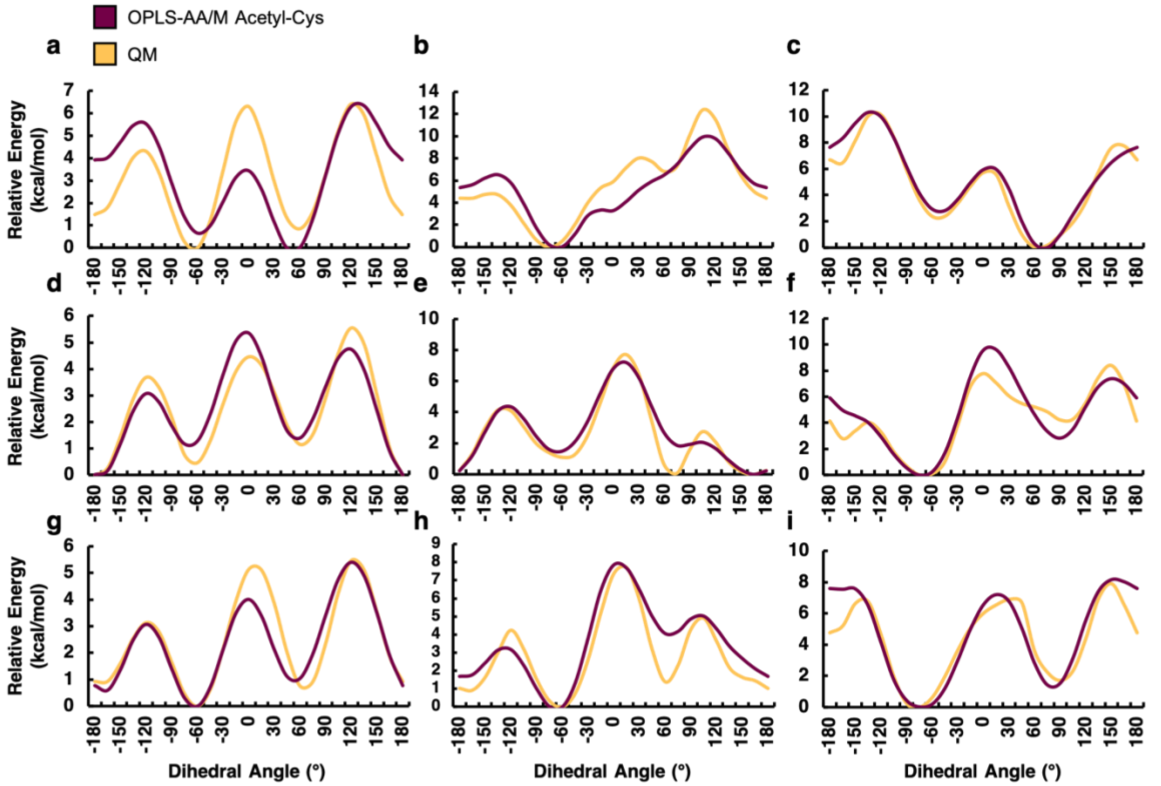


**Figure S4.** Comparison of quantum chemical scans and new OPLS-AA/M force field fits for S-acetyl cysteine in the alpha helical (a,b,c; $\chi_2$= 180, -60, 60), beta sheet (d,e,f; $\chi_2$= 180, -60, 60), and polyproline II (g,h,i; $\chi_2$= 180, -60, 60) conformations.

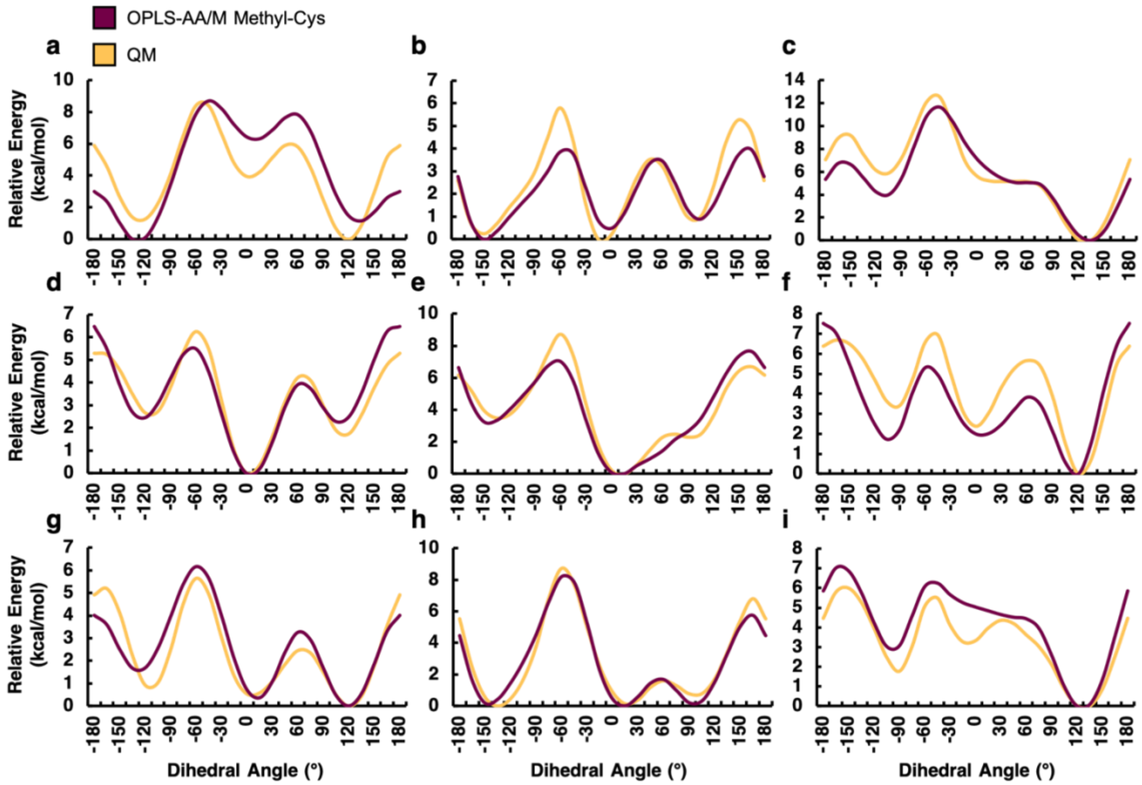


**Figure S5.** Comparison of quantum chemical scans and new OPLS-AA/M force field fits for S-methyl cysteine in the alpha helical (a,b,c; $\chi_2$= 180, -60, 60), beta sheet (d,e,f; $\chi_2$= 180, -60, 60), and polyproline II (g,h,i; $\chi_2$= 180, -60, 60) conformations.

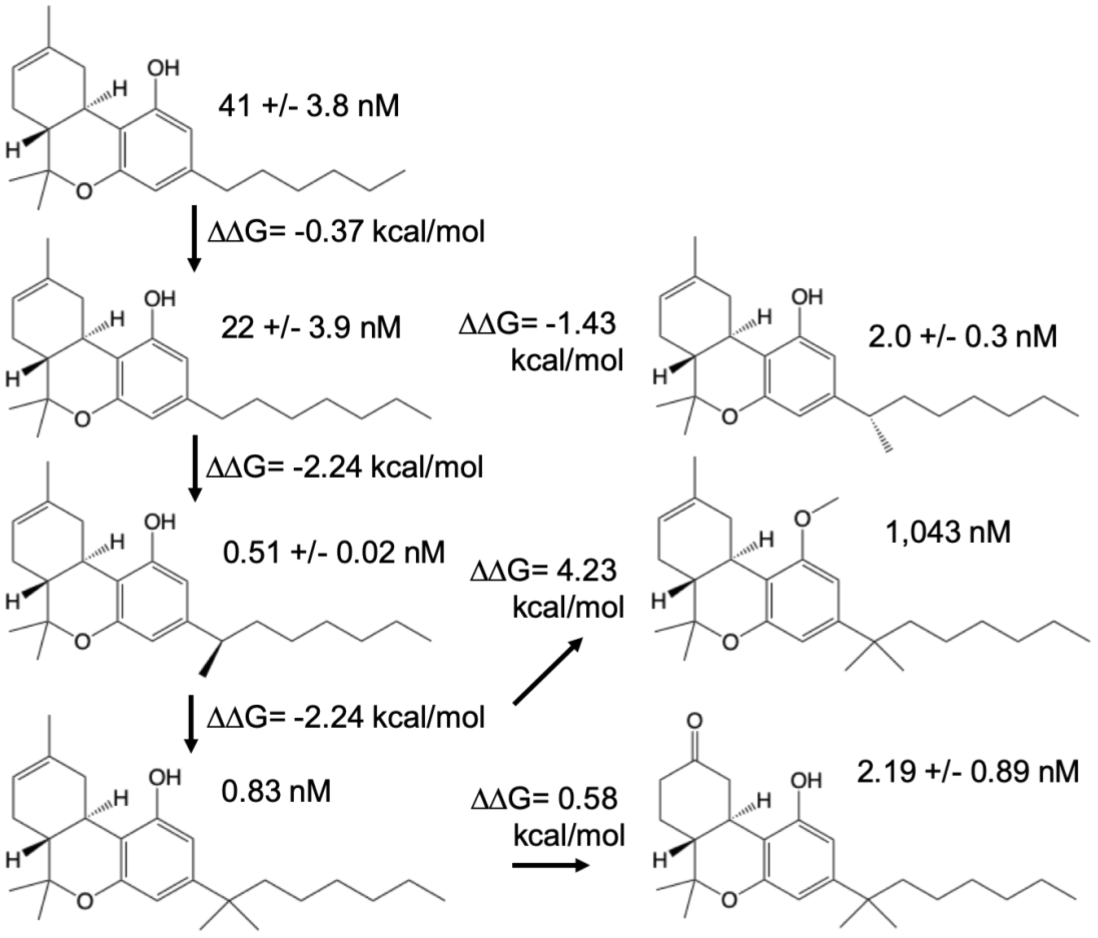


**Figure S6.** Diagram of the Δ8-THC analogues assessed with free energy perturbation simulations with both binding affinities and associated relative free energies of binding.

**Table S3:** Phytocannabinoid relative free energies of binding to CB1 receptor

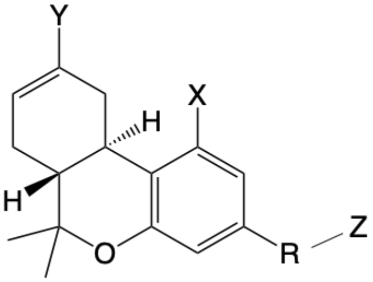


| FEP | R | Z | X | Y | ΔΔG CHARMM (kcal/mol) | ΔΔG OPLS-AA/M (kcal/mol) | ΔΔG Exp (kcal/Mol) |
|---|---|---|---|---|---|---|---|
| 1 | $-CH_2-$ | $C_5 > C_6$ | OH | $CH_3$ | -1.6 | -0.9 | -0.37 |
| 2 | $-CH_2- > -CH(CH_3)-$, L | $C_6$ | OH | $CH_3$ | -1.5 | -0.9 | -1.43 |
| 3 | $-CH_2- > -CH(CH_3)-$, R | $C_6$ | OH | $CH_3$ | -1.4 | -0.9 | -2.24 |
| 4 | $-CH(CH_3)-$, R > $-C(CH_3)(CH_3)-$ | $C_6$ | OH | $CH_3$ | -0.6 | -1.0 | 0.29 |
| 5 | $-C(CH_3)(CH_3)-$ | $C_6$ | OH > $OCH_3$ | $CH_3$ | 2.5 | 2.3 | 4.23 |
| 6 | $-C(CH_3)(CH_3)-$ | $C_6$ | OH | $CH_3$ > O | 1.50 | 1.64 | 0.58 |

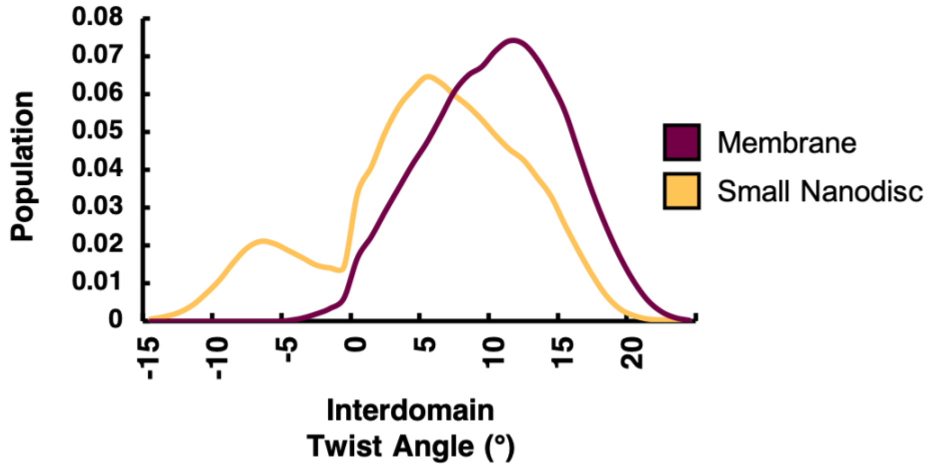


**Figure S7.** Interdomain twist angle of arrestin bound to M2R in either a membrane or a small nanodisc incapable of forming C-edge loop interactions, where higher interdomain twist corresponds to a more 'active-like' conformation of arrestin.

**Table S4:** New torsion parameters reported in this work

| | Dihedral Fourier Coefficient (kcal/mol) | | |
|---|---|---|---|
| Atom Types | V1 | V2 | V3 |
| CL2-CL2-CL2-CL2 | 0.9 | -0.325 | 0.3 |
| CL2-CL2-CL2-CL3 | 1.025 | -0.275 | 0.475 |
| CMc-CMc-CL2-CL2 | -0.2 | 0.425 | -1.875 |
| CMc-CL2-CL2-CL2 | 0.425 | -0.05 | 0.475 |
| CT-CT-OSp-P | -1.42 | -0.62 | 0 |
| N3c-CT-CT-OSp | -3.25 | 0 | 1.5 |
| N3e-CT-CT-OSp | 4.5 | 0.25 | 0.75 |
| CT-OSa-CLL-CL2 | 2 | 9.4 | 0.8 |
| CT-OSa-CLL-OLL | 3.4 | 2.2 | 0.3 |
| OSa-CLL-CL2-CL2 | 0.38 | 0.55 | -0.11 |
| OLL-CLL-CL2-CL2 | -0.545 | 0.33 | 0.015 |
| CLL-CL2-CL2-CL2 | -2.6 | -0.35 | 0.2 |
| CLL-CL2-CL2-CL3 | -2.6 | -0.35 | 0.2 |
| OSp-CT-CT-CT | -2.4 | -0.2 | 2.2 |
| OSp-CT-CT-OSa | 2 | -1.2 | -2.4 |
| OSa-CT-CT-CT | -0.3 | -0.8 | -0.05 |
| OSa-CT-CT-OSa | 4.5 | -2.525 | 0.475 |
| N238-C224-C206-S299 | 2.3 | -0.7 | 1.3 |
| C235-C224-C206-S299 | -2.8 | 0.7 | -0.8 |
| N238-C224-C210-S298 | 2.6 | -0.5 | -0.7 |
| C235-C224-C210-S298 | -2.8 | 0.7 | 1.1 |
| N238-C224-C157-O5447 | 3.7 | -0.9 | 0.3 |
| C235-C224-C157-O5447 | -7.3 | 2.7 | 0.7 |
| N238-C224-C158-O5447 | 3.7 | -0.9 | 0.3 |
| C235-C224-C158-O5447 | -7.3 | 2.7 | 0.7 |
| C224-C206-S299-CLL | -1 | -0.65 | -0.6 |
| C206-S299-CLL-CL2 | 0 | 0 | 0.3535 |
| C206-S299-CLL-OLL | 1.7 | 10.589 | 0 |
| S299-CLL-CL2-CL2 | -0.225 | 0.85 | -1.3 |
